# Modeling interaction between focal cerebral ischemia and Alzheimer’s Disease Pathology in an APP/PS1 Mouse Model

**DOI:** 10.1101/2025.09.05.674491

**Authors:** Paxton K. Waits, P. Anthony Otero, Madison P. Zimmerly, Vidhaath P. Vedati, Kavi J. Shah, James E. Orfila, Paco S. Herson

**Affiliations:** Department of Neurosurgery, The Ohio State University College of Medicine, Columbus, OH, 43210

**Author notes:** Corresponding Author: Paco S. Herson, PhD, Correspondence.

## Abstract

Ischemic stroke is one of the leading causes of death and disability in the United States and is a known risk factor for Alzheimer’s Disease (AD) development. One of the characteristics of AD is the accumulation of β-amyloid peptide due to the proteolysis of Amyloid Precursor Protein (APP) by the protein Presenilin 1 (PS1). The study of interaction between ischemia and AD has been limited due to lack of experimental models. The current study describes a novel model utilizing a brief transient ischemia (15 min MCAO) in an AD mouse model to assess mechanistic interaction. Here we investigate the effects of increased β-amyloid peptide on motor coordination, behavioral and cellular memory, when subjected to brief focal ischemia. APP/PS1 or Wt male mice are initially subjected to either a 15-minute MCAO or Sham surgery. Injury volume using MRI is assessed at 3-days using T2 imaging and demonstrates that injury magnitude was not different between Wild-type and APP/PS1 mice. We observed that 15 min MCAO did not cause significant motor deficits in either WT or APP/PS1 mice, as measured by open field and tapered balance beam. Short-term memory was assessed 7 days after recovery from brief MCAO using the contextual fear conditioning task. APP/PS1 mice exhibited intact memory at 3 months of age and Wt mice also having intact memory 7 days after 15 min MCAO. In contrast, APP/PS1 + 15 min MCAO mice exhibited a significant reduction in freezing behavior suggesting impairment in short term memory. Consistent with our contextual fear conditioning, we observed impaired hippocampal long-term potentiation (LTP) in APP/PS1 + 15 min MCAO compared to their sham counterparts. Remarkably, we observed an additive effect of ischemia and AD, with cellular and behavior memory deficits observed in APP/PS1 mice exposed to brief ischemia that does not cause symptoms in WT mice. This study represents an important new model to be study the mechanistic link between ischemia and accelerated AD progression.

## Introduction

Alzheimer’s disease (AD) is a progressive neurodegenerative disorder characterized by pathological accumulations of amyloid-β (Aβ) and tau proteins, ultimately leading to cognitive decline and functional impairments^1^. While considerable effort has been devoted to understanding the molecular cascades driving AD, the role of cerebral ischemia in modulating the progression of AD pathology remains incompletely defined. The intersection between ischemic events and neurodegeneration has gained prominence as epidemiological data point to a heightened risk of cognitive decline following focal cerebral ischemia^2, 3^. In this study, we address how a small focal ischemic injury, induced by a brief 15-minute middle cerebral artery occlusion (MCAO), may accelerate or exacerbate AD-related abnormalities in an APP/PS1 transgenic mouse model.

To date, most research investigating ischemic stroke in transgenic AD models has focused on more prolonged occlusion times or later disease stages, overlooking how short-duration ischemia could contribute to the early onset of AD phenotypes. By building on existing literature that demonstrates the vulnerability of AD brains to vascular insults, this work explores whether mild ischemia significantly alters behavioral and pathological outcomes compared to wild-type (WT) controls. While some prior studies suggest that AD-transgenic mice exhibit worse stroke outcomes than non-transgenic counterparts^4, 5^, how mild ischemia, particularly in younger animals, remain poorly understood. Addressing this gap can provide novel insight into how transient vascular events may influence the trajectory of amyloid-driven neuropathology.

Accordingly, this work examines the effects of short-duration MCAO (15-min) on motor coordination, contextual fear conditioning, and hippocampal plasticity in APP/PS1 mice. Our hypothesis is that brief ischemia would exacerbate and potentially accelerate AD-like pathology, reflecting a vulnerability that might otherwise go unrecognized in early disease stages. By integrating MRI-based lesion analysis with behavioral and electrophysiological measures, our methodology provides a multifaceted perspective on how vascular and amyloid pathologies converge. In doing so, this research aims to characterize an experimental model to be utilized in future studies to shed light on potential mechanisms underlying vascular contributions to AD progression, ultimately guiding future therapeutic strategies that address both ischemic and neurodegenerative processes.

## Results

### T2 MRI Analysis Following MCAO

MRI analysis using T2-weighted imaging revealed distinct patterns of ischemic injury following 15-minute middle cerebral artery occlusion (MCAO) in both APP/PS1 transgenic and WT mice. Hyperintense lesions were observed within the ipsilateral hemisphere of APP/PS1 and WT mice, indicating cerebral edema and tissue damage. Quantitative assessment of lesion volume demonstrated a substantial increase in injury severity in mice post-MCAO. No significant difference was found between the APP/PS1 MCAO and WT MCAO groups (17.1 ± 5.3%; n=7 and 21.2 ± 6.7%; n=12, respectively, p>0.05, unpaired t-test) (Figure 1).

**Figure 1:**
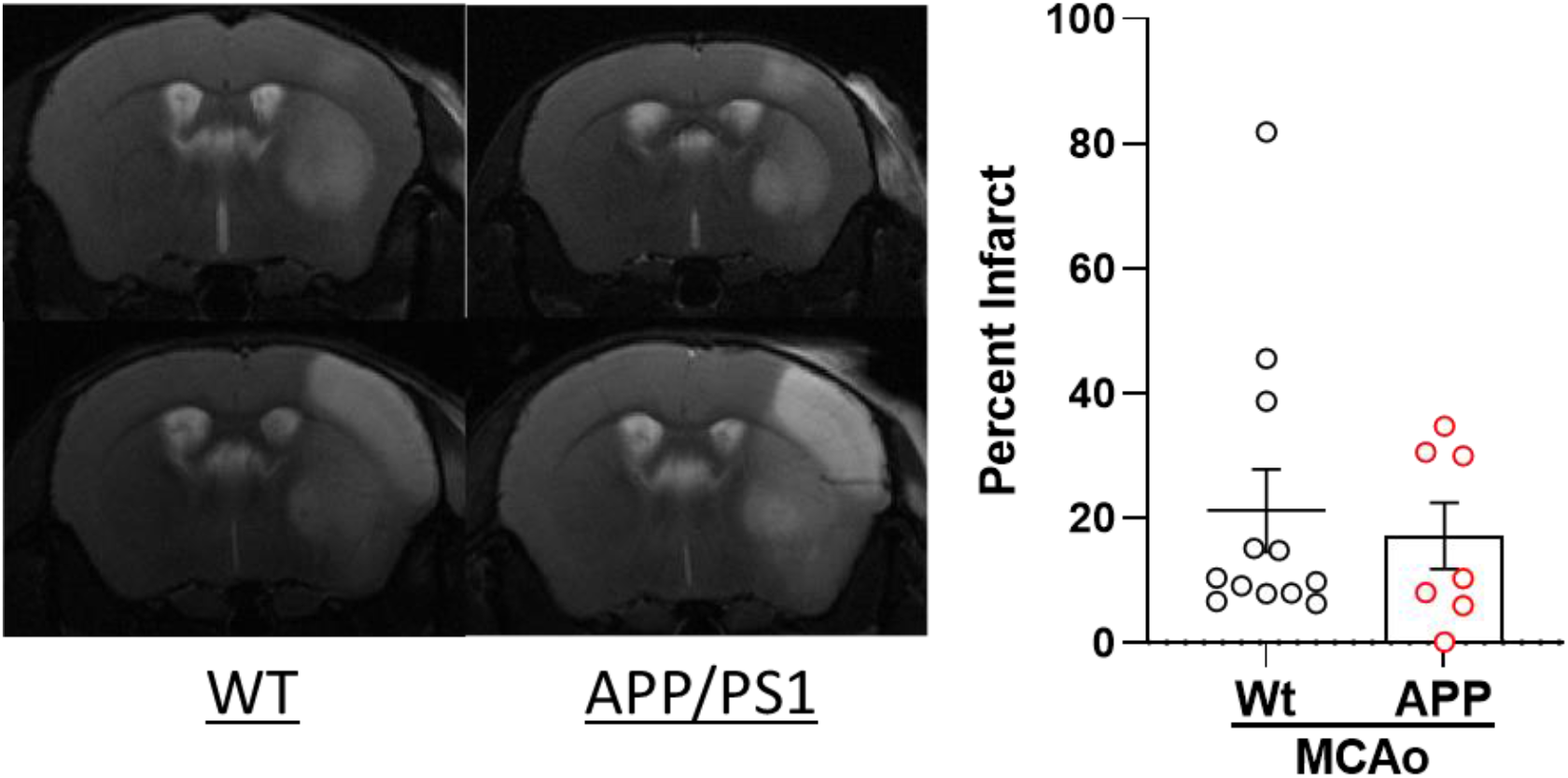
APP/PS1 transgenic mice do not exhibit increased infarct volume following 15-minute MCAO. Images were obtained 3-days following MCAO. Each point represents one animal. A) Images of a mouse brain at the level of the striatum were taken from the two animals closest to the mean. B) Quantification of the percent infarct of the ipsilateral hemisphere. (n = 7 and 12 for APP and Wt, respectively, p>0.05, unpaired t-test).

### Impact of 15-min MCAO on Motor Coordination

Motor coordination was assessed via tapered balance beam performance, quantified by counting total slips as well as categorized slips (front, back, right, and left slips). It was observed that 15-minute MCAO injury did not result in an increase in total foot slips associated with motor deficits in WT mice (8.3 ± 4.6 slips, n=8) when compared to WT sham mice (3.1 ± 1.23 slips, n=8). Similarly, no significant motor deficits were observed in APP/PS1 MCAO mice (10.4 ± 3.8 slips, n=8) when compared to APP/PS1 sham mice (4.7 ± 1.8 slips, n=7) (Figure 2). These findings suggest that APP/PS1 mice do not exacerbate the effects of 15-minute MCAO on motor coordination.

**Figure 2.**
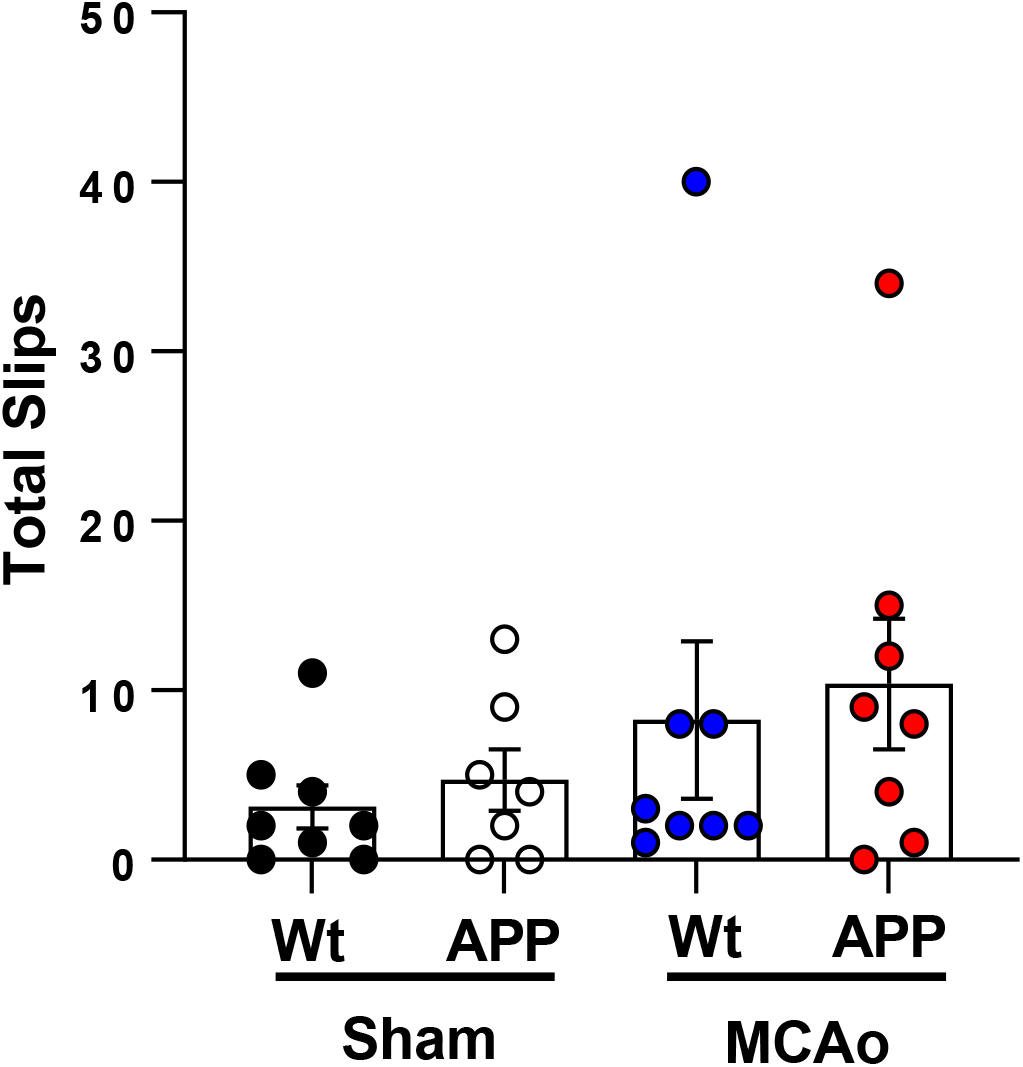
A 15-minute middle cerebral artery occlusion (MCAO) does not impair motor coordination. Deficits following MCAO were assessed by counting total number of slips in wild-type (WT) and amyloid precursor protein (APP) transgenic mice. Neither WT MCAO (blue filled circles) nor APP/PS1 (red filled circles) mice showed a significant difference in total foot slips when compared to WT sham (black filled circles) or APP/PS1 sham (open circles) mice. Quantification of total foot slips from individual mice were analyzed and scored by a blinded observer. (n = 7-8/group. p > 0.05, 1-Way ANOVA).

### Contextual Fear Conditioning Performance Post-MCAO

To determine whether a 15-minute ischemic event affected memory, we performed contextual fear conditioning 7 days after MCAO. Contextual fear memory was evaluated by measuring the percentage of freezing behavior during the conditioning (Day 1) and recall (Day 2) phases. Figure 3d shows that 15-minute MCAO injury failed to cause memory deficits in WT mice (41.7 ± 9.2%, n=8) when compared to WT Sham (50.0 ± 7.7%, n=8). In contrast, a significant decrease in freezing behavior associated with memory function was observed in APP MCAO-treated mice (26.3 ± 6.2%, n=8) compared to APP/PS1 sham mice (57.5 ± 8.3%, n=8). These findings suggest that the impaired memory observed may be attributed to the early vulnerability of APP/PS1 transgenic mice to MCAO-induced cognitive deficits.

**Figure 3.**
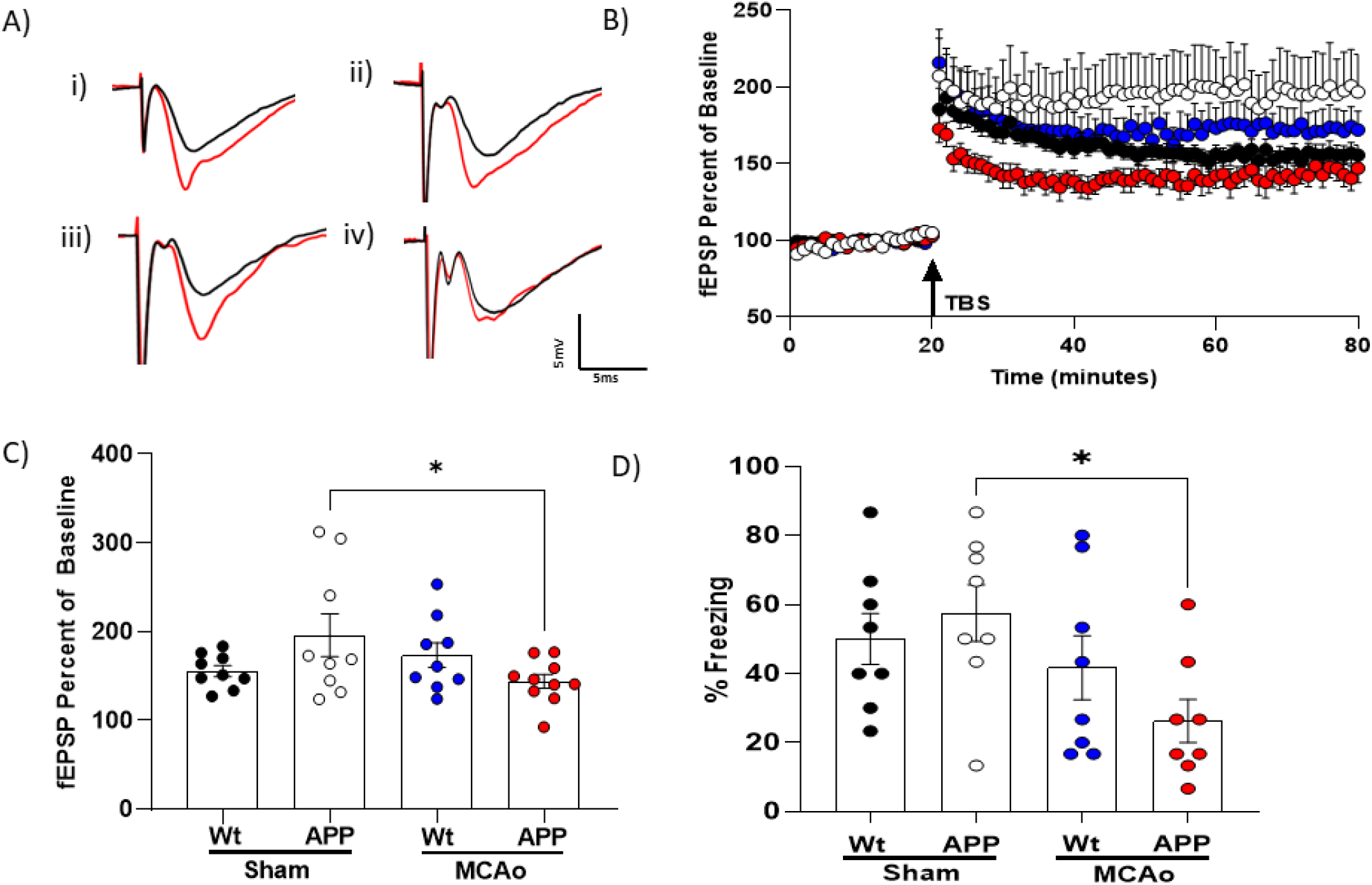
APP transgenic mice exhibit impaired hippocampal plasticity and memory function 7 days after middle cerebral artery occlusion (MCAO). (A) Representative fEPSP traces from (i) WT sham, (ii) APP Sham, (iii) WT MCAO, and (iv) APP MCAO mice at 7 days post-surgery. B) Time course of fEPSP slope 7 days after surgery in WT Sham (black filled circles), APP Sham (open circles), WT MCAO (blue filled circles) and APP MCAO (red filled circles). The arrow indicates the timing of TBS (40 pulses). C) Quantification of change in fEPSP slope after 60 min following TBS, normalized to baseline, set at 100%. Each point represents a hippocampal slice that was recorded with no more than two slices per animal. D) Memory impairment 7 days after MCAO. APP MCAO-injured mice displayed significant memory dysfunction after contextual fear conditioning 7 days after injury. Quantification of freezing behavior scored 24 h after contextual fear conditioning in a novel environment (n = 8/group. *p < 0.05 compared to sham, 1-Way ANOVA).

### Long-Term Potentiation Impairment Following MCAO

To assess the effect on hippocampal long-term potentiation (LTP), extracellular field recordings of CA1 neurons were collected from acute hippocampal slices prepared 7 days after MCAO. Field excitatory postsynaptic potentials (fEPSPs) were elicited by stimulating the Schaffer collaterals and recorded in the stratum radiatum of CA1. After establishing a 20-minute stable baseline, LTP was induced using a 40-pulse theta-burst stimulation (TBS), as described in the methods, and recorded for 60 minutes post-TBS. Figure 3 B, C shows that a 15-minute MCAO injury failed to impair LTP in WT MCAO mice (173.6 ± 13.9%, n=9) 7 days after MCAO when compared to WT sham mice (155.6 ± 6.4%, n=9). In contrast, a significant LTP deficit was observed in the APP/PS1 MCAO mice (143.9 ± 7.8%, n = 10) 7 days after MCAO, when compared to the APP/PS1 sham mice (196.0 ± 24.1%, n = 9). This data aligns with our behavioral findings and suggests that the adverse effects of MCAO on hippocampal synaptic function may be linked to the progression of AD pathology.

## Discussion

The present study sought to establish whether mild ischemic events could exacerbate or accelerate AD onset. We demonstrate that APP/PS1 transgenic mice exhibit increased vulnerability to ischemic injury following a brief 15-minute MCAO, as evidenced by pronounced cognitive deficits. The impaired performance of APP/PS1 mice seen in contextual fear conditioning and impaired hippocampal long-term potentiation (LTP), underscores the exacerbation of ischemia-induced neurological dysfunction in the presence of amyloid pathology. Importantly, this additive negative effect was observed independent of alterations in injury magnitude, demonstrating a complex interaction of Aβ and ischemia on hippocampal function.

To the best of our knowledge, this is the first report evaluating the impact of a brief ischemia (sub-symptomatic MCAO) in young (8–12 weeks old) APP/PS1 mice, thereby expanding on existing short-duration stroke models in transgenic animals. In line with previous findings, our T2-weighted MRI results confirm that even a brief occlusion can induce robust hyperintense abnormalities^6^. Moreover, other studies have shown that both Alzheimer’s disease (AD) animal models^4, 7^ and AD patient populations exhibit increased susceptibility to ischemic injury, paralleling our observation that APP/PS1 mice are more sensitive to stroke than their wild-type counterparts. A particular strength of this work is the comprehensive assessment of both motor and cognitive domains alongside MRI, which offers a multidimensional view of ischemic outcomes. Consistent with previous studies, we observed intact short-term memory and hippocampal plasticity in young APP/PS1 mice^8^. Remarkably, we observed significant cellular and behavioral memory deficits in young APP/PS1 mice exposed to a brief ischemic insult (15 min) that failed to cause behavioral or cellular deficits in young wild-type mice. This data is consistent with our hypothesis that cerebral ischemia increases the magnitude of Aβ-induced brain dysfunction and likely accelerated AD progression. The observed exacerbation of cognitive impairments in APP/PS1 mice suggests that even a transient vascular insult could trigger earlier or more severe disease progression. This work underlines the importance of vascular contributions to AD and demonstrates how brief episodes of ischemia may serve as a catalyst for pathological processes in at-risk populations. Although this data strengthens the case for integrating cerebrovascular considerations into AD research, additional studies are necessary to delineate the underlying mechanisms. Specifically, it will be important to assess cellular responses (microgliosis, astrogliosis) to ischemia in this model as well as how Aβ deposition, and tau pathology interact over longer time frames. Future research should also explore whether mitigating inflammatory or vascular damage after mild ischemia might slow disease progression in AD models, thereby providing new avenues for therapeutic intervention.

## Methods

### Tapered Balance Beam

Motor coordination was assessed using the tapered balance beam test^9, 10^. The apparatus consisted of two platforms: a starting platform at the base and an enclosed, minimally lit box at the summit. A tapered beam connected the two platforms, becoming narrower toward the summit, with a translucent plastic ledge positioned beneath the beam to catch mice in the event of slips. During both learning and recording phases, mice were transported in white buckets.

On day 7 following MCAO, mice were first allowed to acclimate to the enclosed box for 2 minutes. After acclimation, the learning phase began, consisting of three separate runs starting sequentially from ¾, ½, and ¼ distances up the beam. Immediately following the learning phase, the mice completed two recorded full-length crossings of the beam. Later, a blinded observer analyzed the first video recording to quantify slips, defined as any step made onto the translucent plastic ledge rather than the beam itself.

### Contextual fear conditioning

Contextual fear conditioning was employed as a hippocampal-dependent memory assessment, following previously established paradigms^11, 12^. The apparatus consisted of two fear-conditioning chambers with shock-grid floors, each composed of 16 stainless steel rods connected to a shock generator (Colbourn Instruments, Model H13-15, Whitehall, PA, USA). Mice were transported to and from testing sessions in white buckets. On day 7 post-MCAO (immediately following the balance beam test), mice underwent two separate 2-minute habituation sessions in the conditioning chamber. After the second habituation session, mice received a single foot shock (2 seconds, 1.0 mA). Following the shock, animals were returned to their home cages and transported back to the vivarium. Testing occurred 24 hours later, wherein mice were placed back into the same conditioning chambers. Freezing behavior—defined as the absence of movement aside from heartbeat and respiration—was measured in 10-second intervals over a 5-minute test by a blinded observer.

### Hippocampal Slice Preparation

Hippocampal slices were prepared 7 days post-MCAO, as previously described^11-14^. Mice were anesthetized with 3% isoflurane in an oxygen-enriched chamber, perfused with artificial cerebrospinal fluid (aCSF) (consisting of in mmol/L: 125 NaCl, 2.5 KCl, 25 NaHCO_3_, 1.25 NaH_2_PO_4_, 2.0 CaCl_2_, 1.0 MgCl_2_, and 12 glucose), and their brains rapidly extracted. Extracted brains were placed in ice-cold (2–5°C), oxygenated (95% O_2_/5% CO_2_) aCSF. Horizontal hippocampal slices (300 µm thick) were cut using a Vibratome 1000 (Leica) and subsequently transferred to a holding chamber filled with oxygenated aCSF, incubated for 1.5–2 hours prior to electrophysiological recording.

### Electrophysiology

Field excitatory postsynaptic potentials (fEPSPs) were recorded from hippocampal slices, as previously described^11-14^. Slices were placed in a temperature-controlled (31 ± 0.5°C) interface chamber, continuously perfused with oxygenated aCSF at a rate of 2.0 mL/min. Synaptic field potentials were elicited by stimulating the Schaffer collaterals and recorded in the CA1 hippocampal region, adjusting stimulation intensity to achieve 50% of the maximum fEPSP slope. Paired-pulse stimulation utilized a 50 ms interpulse interval (20 Hz), with responses quantified as the ratio of the second pulse to the first.

Baseline responses were collected at 20-second intervals over 20 minutes at 50% of the maximum fEPSP slope. After baseline stabilization, theta burst stimulation (TBS)— consisting of ten bursts of four pulses at 100 Hz delivered every 200 ms—was administered. Following TBS, fEPSP responses continued to be recorded at 20-second intervals for 60 minutes. Analog fEPSPs were amplified (1000×), filtered through a Grass Instruments pre-amplifier (Model LP511 AC) at 1.0 kHz, and digitized at 10 kHz. To quantify synaptic potentiation, fEPSP slopes from the final 10 minutes of recording (50– 60 minutes post-TBS) were averaged and normalized to the last 10-minute baseline segment. Data were represented as percent change from baseline. Each data point corresponded to a single hippocampal slice, with no more than two slices per animal to ensure biological variability.

### T2 MRI

Infarct volumes were measured using T2 MRI imaging conducted 3 days post-MCAO. Mice were anesthetized initially with 3% isoflurane, then transferred to the MRI system and maintained on 1% isoflurane for a total duration of 15 minutes. MRI images, captured every 0.5 mm along the cerebral cortex, were analyzed by a blinded observer using ITK-SNAP software. Data were expressed as infarct volumes, quantified as a percentage of the ipsilateral hemisphere.

### 15-min MCAO

Focal cerebral ischemia was induced using an intraluminal filament, as previously described^11, 15-17^. Mice were induced using 95% oxygen and 5% carbon dioxide at 300 mL/min or greater with 5% or less isoflurane. Maintenance was achieved using the same oxygen and carbon dioxide parameters as induction with 97-103 mL/min of airflow with 2% or less isoflurane. A temporary suture was tied onto the External Carotid Artery and a removable aneurism clip was used on the Common Carotid to avoid blood loss. Ischemia was induced by entering the Internal Carotid Artery and advancing the intraluminal filament to occlude the Middle Cerebral Artery. Occlusion was confirmed using a laser Doppler probe (Moor Instruments VMS-LDF2) that was located over the ipsilateral parietal cortex. Occlusion was considered a blood flow deduction lower than 25% of the baseline value. Once the occlusion was confirmed, a suture would be tied around the Internal Carotid Artery and the filament, followed by the removal of the suture around the External Carotid Artery and the aneurism clip on the Common Carotid. Once 15 minutes had passed from the time the intraluminal filament was placed, the filament would be removed, anesthesia delivery would be stopped, and the mouse would return to an individual cage to recover. Sham animals were exposed to anesthesia and underwent arterial access without any occlusion.

### Experimental animals

All experimental protocols were approved by The Ohio State University College of Medicine Institutional Animal Care and Use Committee and conducted in accordance with National Institutes of Health guidelines for animal research. A total of 69 adult male APP/PS1 and C57Bl/6 mice (20–25 g, 8–12 weeks old) from our colony—originally obtained from Jackson and Charles River Laboratories, respectively—were utilized.

Animals were housed under standard 12-hour light-dark cycles with free access to food and water. All experiments adhered to ARRIVE guidelines^18^, and analyses were performed by blinded individuals. Experimental groups included MCAO and sham-treated APP/PS1 and WT mice, with survival periods of 7 days for behavioral tests (n=7-8 mice/group) and electrophysiology experiments (n=6–7 mice/group).

### Statistical Analysis

All data presented as mean±SEM. Statistical analysis of all data was determined using the Student’s t-test for two group comparisons and one-way analysis of variance (ANOVA) and post-hoc Dunnett’s test for comparison of multiple groups. Differences considered statistically significant with P<0.05.

